# From Network Topology to Parameterization: An Optimized Model of Endothelial Mechanobiology

**DOI:** 10.64898/2026.10.05.756772

**Authors:** Pete H. Gueldner, Linda Irons, Jeremy L. Warren, David S. Li, Jay D. Humphrey

## Abstract

Endothelial cells integrate biomechanical and biochemical cues to regulate vascular function, which can be modeled using signaling networks. Yet, the extent to which their signaling behavior is determined by network topology versus reaction-specific parameterization remains unclear. We developed a continuous logic-based ordinary differential equation model of endothelial signaling containing 70 species and 123 regulatory reactions assembled from the literature. Model predictions were evaluated against 79 literature-defined responses to imposed wall stress, wall shear stress, hypoxia, or TNFα. We first tested whether a single global parameter triplet (describing reaction weight, Hill coefficient, and half-maximal activation) could reproduce these observations. Investigator-selected global parameters reproduced 62/79 responses (78.48%). Monte Carlo sampling of 100,000 globally constant parameter sets improved agreement to 69/79 responses (87.34%), indicating that network topology alone captured most qualitative signaling behavior but remained sensitive to parameter choice. Allowing reaction-specific parameters, also by Monte Carlo sampling within 100,000 additional parameter sets, increased the best agreement to 75/79 responses (94.94%). We then used a genetic algorithm to identify the smallest subset of reactions requiring reaction-specific specification while retaining a common constant-parameter spine for the remainder of the network. Near complete agreement with literature-defined responses (78/79, or 98.73%) was achieved with reaction-specific parameterization of only 18 of 123 reactions. The optimized model was subsequently used to simulate responses to continuous changes in wall shear stress, revealing pathway-dependent nonlinear responses including monotonic increases in nitric oxide and pSmad1/5, nonmonotonic responses in endothelin-1 and NF-κB, and a biphasic pSmad2/3 response. These findings show that most observed endothelial signaling behavior can be reproduced by a fixed network topology and broadly shared reaction parameters with only a limited subset of interactions requiring quantitative specialization. This framework provides a compact endothelial signaling model for investigating mechanobiological regulation and for future integration with multiscale models of vascular adaptation and disease.

**Author Summary:** Endothelial cells line blood vessels and continually respond to changes in hemodynamics and inflammation, helping to maintain vascular health. Understanding how these signals interact is challenging because the strengths of individual connections within cellular signaling networks are often unknown. We developed a computational model of these interactions and investigated whether its behavior was determined primarily by how the network was connected or by the numerical values assigned to individual connections. Using Monte Carlo methods, which repeatedly test randomly selected values, we evaluated 200,000 different parameter combinations. We found that assigning the same values throughout the network reproduced approximately 87% of reported experimental observations, whereas allowing values to differ between reactions increased agreement to 95%. We then combined Monte Carlo sampling with an additional optimization method to identify connections that required individual adjustment. Remarkably, modifying only 18 of 123 connections increased agreement to 98%. Our findings demonstrate how Monte Carlo methods can help distinguish the importance of network structure from individual signaling interactions, providing a systematic approach for developing simpler, more reliable models of cellular behavior and vascular disease.

## Introduction

The endothelium consists of a contiguous monolayer of endothelial cells and performs many vital functions that ensure vascular health (1, 2). It serves, for example, as a selective permeability barrier between the flowing blood and mural contents; it plays a critical role in hemostasis; and it plays a key role in immune cell regulation. From the perspective of biomechanics, the endothelium also serves as a vital mechanosensor of local blood flow and vascular distension, which allows it to influence smooth muscle cell vasoactivity via a mechano-regulated control of the synthesis and release of dilators (nitric oxide - NO) and constrictors (endothelin-1 - ET1). Recent advances in modeling vascular mechanobiology have included development of intracellular cell signaling models for vascular smooth muscle cells (3), pulmonary arterial fibroblasts (4), and brain endothelial cells (5). The latter integrated multiple cerebrovascular endothelial signaling pathways to investigate disease-associated signaling, but a corresponding model focused on arterial endothelial mechanobiology and its responses to mechanical, inflammatory, and vasoactive stimuli remains lacking.

Intracellular signaling is driven by diverse stimuli, including mechanobiological and immunobiological, and it regulates cell-cell and cell-matrix interactions among many other biological processes. Computational models of cell signaling networks provide a way to formalize pathway knowledge, simulate responses to perturbations, and generate experimentally testable hypotheses (6, 7). Many signaling models begin as directed regulatory graphs wherein nodes represent signaling species or phenotypic markers and edges represent activating or inhibitory interactions (8–10). This structure is attractive because it can be assembled from observations reported via in vitro and in vivo studies, even when complete kinetic measurements are unavailable (11, 12). Recent benchmarking of automatically generated signaling networks demonstrated that incomplete reconstruction of literature-curated network topologies associates with poor predictions of experimentally observed perturbation results, emphasizing the importance of accurate network construction and functional validation (13).

Although network topology can often be defined from prior biological knowledge, the parameters that govern edge strength and activation sensitivity are seldom known for the many chemical species-to-species interactions. In signaling models based on Hill functions, parameters such as reaction weight (*w*), Hill coefficient (*n*), and half-maximal activation (*EC_50_*) determine whether a regulatory edge acts weakly or strongly, gradually or switch-like, and at low or high upstream activity (14, 15). In practice, large literature-derived signaling models are often initialized using manually selected or globally constant parameter values when direct measurements are unavailable (3, 4, 11). Parameter uncertainty has been examined in prior logic-based vascular signaling models using sampling-based uncertainty quantification (4). This approach can be useful for testing whether network topology alone produces reasonable qualitative behavior, but it does not determine whether the selected values are unique, robust, or optimal. Monte Carlo sampling provides a practical way to explore a large parameter space, identify experimentally consistent parameter sets, and quantify whether acceptable model behavior is limited to a narrow region or distributed across many parameter combinations (16–18).

A key unresolved question is whether experimentally observed signaling phenotypes are encoded primarily by network topology or whether they require reaction-specific parameterization. If topology is sufficient, a single global set of *w*, *n*, and *EC_50_* values should reproduce directional (i.e., increases versus decreases) experimental trends across perturbations; if not, improved agreement should require some level of group-or reaction-specific parameter variation, suggesting that particular signaling modules require distinct strengths, sensitivities, or nonlinearities. The goal of the present work is two-fold: to develop an arterial endothelial cell signaling model and to demonstrate the use of Monte Carlo simulations to determine when the signaling network topology is sufficient to reproduce experimental phenotypes versus when reaction-specific parameterization is needed. We illustrate the utility of the modeling by comparing investigator-estimated parameters, globally constant Monte Carlo-selected parameters, and perturbed reaction-specific parameter sets across experimentally defined conditions relevant to endothelial cell mechanobiology.

## Results

### Endothelial intracellular signaling network architecture

The endothelial intracellular signaling network was constructed to integrate biomechanical and biochemical stimuli through receptor-level signaling, intracellular intermediates, transcriptional regulators, and phenotypic outputs (**Fig. 1**). The resulting network contained 70 species and 123 reactions. Mechanical and extracellular inputs included stress, WSS, hypoxia, TNFα, and AngII, with downstream signaling mediated through pathways including integrins, ion channels, VEGFR, TGFβ receptors, TNFαR, endothelin receptors, and AngII receptors. These pathways converged on signaling intermediates and transcriptional regulators including PI3K/Akt/mTOR, MAPK signaling, RhoA/ROCK, Smads, NF-κB, HIF1α, and Klf2, and ultimately regulated vasoactive mediators and phenotypic outputs including NO, ET1, migration, proliferation, apoptosis, and EndoMT.

**Figure 1.**
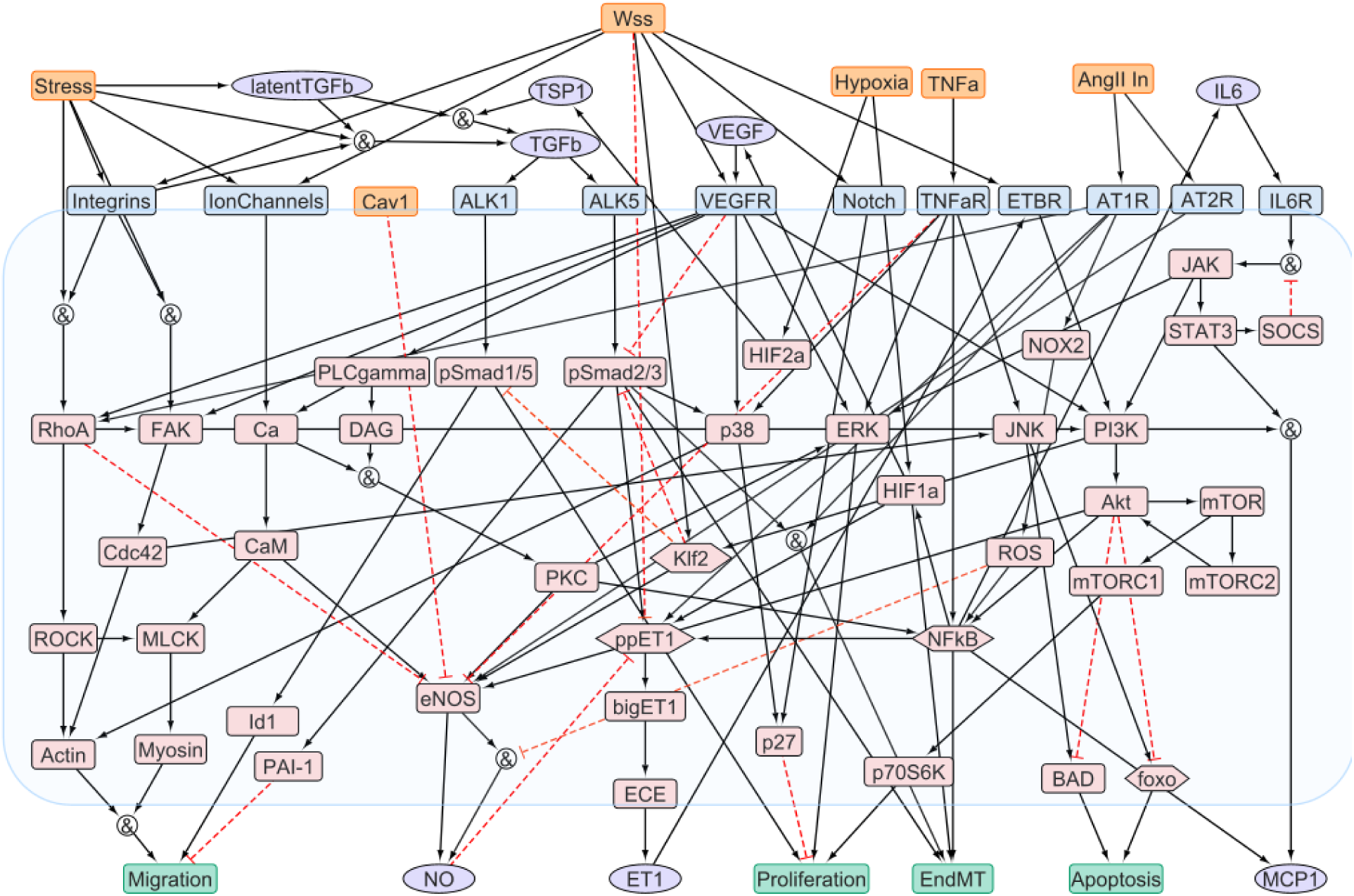
Endothelial intracellular signaling network. Logic-based ordinary differential equation model of a representative endothelial cell integrating biomechanical and biochemical stimuli, including inflammatory. The network contains 70 species and 123 regulatory reactions. Solid black arrows denote activating interactions, red dashed lines denote inhibitory interactions, and nodes containing an ampersand denote AND relationships betwee upstream signals

Model predictions were evaluated against a literature-derived validation matrix containing 79 directional comparisons across independent perturbations in stress, WSS, hypoxia, and TNFα. Agreement was assessed according to whether the predicted change in each species was consistent qualitatively with the experimentally reported increase, decrease, or lack of change.

### Investigator-selected parameters reproduce most literature-defined signaling responses

The initial model used investigator-selected global reaction parameters of *w*=0.85, *n*=1.4, and *EC_50_*=0.6 for reactions without separately prescribed values. This investigator-tuned parameterization correctly reproduced 62 of the 79 literature-defined responses (**Table 1**), corresponding to 78.48% agreement with the validation matrix (**Fig. 2**). Thus, the literature-derived network architecture coupled with a single manually selected parameter set was sufficient to reproduce directional changes for most of experimentally reported responses.

**Figure 2.**
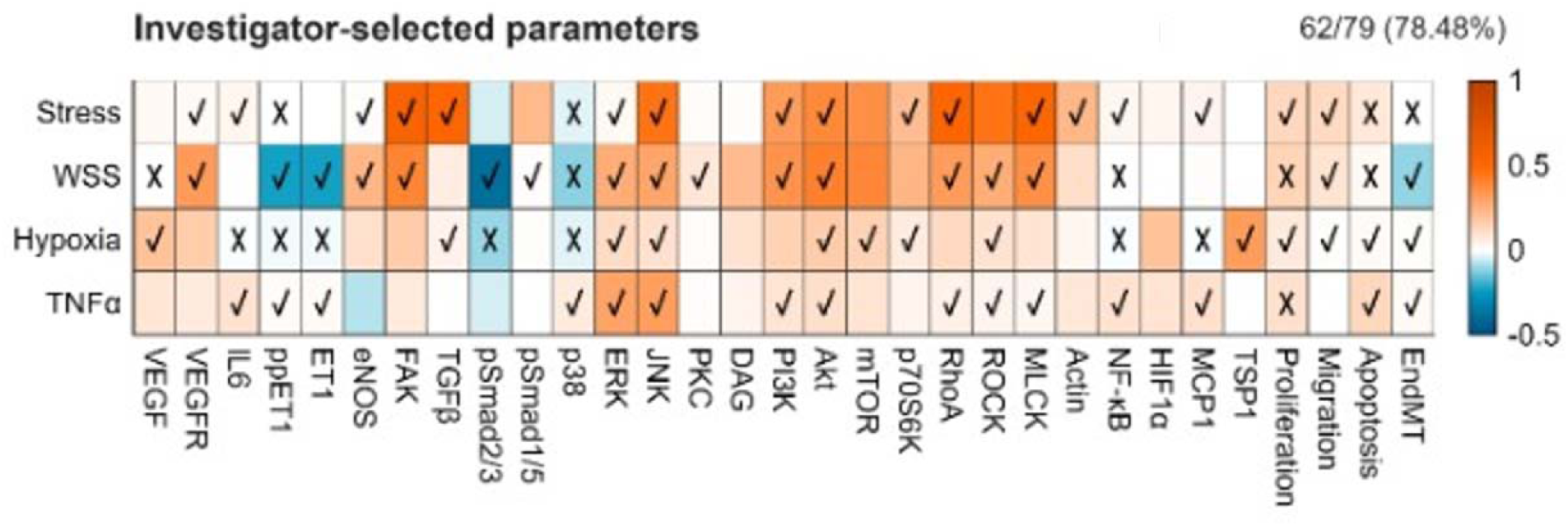
Validation of the investigator-selected parameterization against literature-defined endothelial responses. Heatmap showing model-predicted changes in signaling species and phenotypic outputs. Backgroun color indicates the simulated change in normalized activity relative to the corresponding baseline state, with orang indicating an increase, blue a decrease, and white little or no change. Using investigator-selected parameters (*w*=0.85, *n*=1.4, and *EC_50_*=0.6) for reactions without separately prescribed values, the model correctly reproduced 6 of 79 literature-defined responses (78.48%).

**Table 1.** Literature-derived validation matrix of endothelial cell signaling and phenotypic responses to biomechanical and biochemical stimuli. Orange indicates upregulated, blue italic indicates downregulated, black hyphen indicates no change. Abbreviations: HUVEC, human umbilical vein endothelial cells; HAEC, human aortic endothelial cells; BAEC, bovine aortic endothelial cells; RAEC, rat aortic endothelial cells; HPAEC, human pulmonary artery endothelial cells; BPAEC, bovine pulmonary artery endothelial cells; PMVEC, pulmonary microvascular endothelial cells; HSaVEC, human saphenous vein endothelial cells; HMEC, human microvascular endothelial cells; VEC, valvular endothelial cells. Literature citations given as (#).

| **Species** | **Stress** | **WSS** | **Hypoxia** | **TNFα** |
|---|---|---|---|---|
| **VEGF** | — | **(19) HUVEC ↥** | **(20) BPAEC ↥** | — |
| **VEGFR** | **(21) HUVEC ↥** | **(19, 22) HUVEC HAEC ↥** | — | **(23) HUVEC ↥**<br>***(24) HUVEC, HAEC ↓*** |
| **IL6** | **(25) HUVEC ↥** | — | **(26) HUVEC ↥** | **(27, 28) HUVEC ↥** |
| **ppET1** | **(29) HUVEC ↥** | ***(29) HUVEC ↓*** | **(30) HUVEC ↥** | **(31) BAEC ↥** |
| **ET1** | — | ***(32) HUVEC ↓*** | **(30) HUVEC ↥** | **(31) BAEC ↥** |
| **eNOS** | **(33, 34) HUVEC, BAEC ↥** | **(35) BAEC ↥** | ***(36-38) HSaVEC, BPAEC, HUVEC ↓***<br>**(39) BAEC ↥** | **(40) HUVEC, HMEC ↥**<br>***(41, 42) BAEC, HUVEC ↓*** |
| **FAK** | **(43) BPAEC ↥** | **(44, 45) HUVEC, BAEC ↥** | — | — |
| **TGFβ** | **(46, 47) HUVEC, HPAEC ↥** | **(48) BAEC ↥**<br>***(22) HAEC ↓*** | **(49) HUVEC ↥** | — |
| **pSmad2/3** | — | ***(50) HUVEC ↓*** | **(51) HUVEC ↥** | — |
| **pSmad1/5** | — | **(52) HUVEC, HAEC, BAEC ↥** | — | — |
| **p38** | **(47, 53) HPAEC, BAEC ↥** | **(54) BPAEC, HPAEC ↥** | **(55) BAEC ↥** | **(56, 57) HUVEC ↥** |
| **ERK** | **(33, 47, 53) HUVEC, HPAEC, BAEC ↥** | **(44, 58) HUVEC, BAEC ↥** | **(59) PMVEC ↥** | **(56, 60) HUVEC, HMEC ↥**<br>**(61) BAEC -** |
| **JNK** | **(53) BAEC ↥** | **(58) BAEC ↥** | **(55) BAEC ↥** | **(56, 57) HUVEC ↥** |
| **PKC** | — | **(62) HUVEC ↥** | — | — |
| **DAG** | — | — | — | — |
| **PI3K** | **(34, 63) BAEC ↥** | **(64) HUVEC ↥** | — | **(57, 65) HUVEC ↥** |
| **Akt** | **(33, 34, 66) HUVEC, BAEC ↥** | **(35, 64) BAEC, HUVEC ↥** | **(67) RAEC ↥** | **(57, 65) HUVEC ↥** |
| **mTOR** | — | — | **(67) RAEC ↥** | — |
| **P70s6k** | **(68) BAEC ↥** | — | **(67) RAEC ↥** | — |
| **RhoA** | **(47) HPAEC ↥** | **(69) HUVEC ↥** | — | **(70) HUVEC ↥** |
| **ROCK** | — | **(71) BAEC ↥** | **(36) BPAEC ↥** | **(70) HUVEC ↥** |
| **MLCK** | **(47) HPAEC ↥** | **(72) BAEC ↥** | — | **(70) HUVEC ↥** |
| **Actin** | **(73) HUVEC ↥** | — | — | — |
| **NF-κB** | **(25) HUVEC ↥** | **(74) BAEC ↥** | **(75) PMVEC ↥** | **(27, 76) HUVEC, HMEC ↥** |
| **HIF1α** | — | — | — | — |
| **MCP1** | **(77, 78) HUVEC ↥** | — | **(79) HUVEC ↥** | **(76, 80) HMEC, HUVEC ↥** |
| **TSP1** | — | — | **(81) BAEC ↥** | — |
| **Proliferation** | **(68) BAEC ↥** | ***(82) HUVEC, BAEC ↓*** | **(83) HPAEC ↥** | ***(84, 85) BAEC, HAEC ↓*** |
| **Migration** | **(86) BAEC ↥** | **(45) BAEC ↥** | **(87) HUVEC ↥** | — |
| **Apoptosis** | ***(66) BAEC ↓*** | ***(19, 64) HUVEC ↓*** | **(88) HAEC ↥** | **(85) HAEC ↥** |
| **EndMT** | **(89, 90) HUVEC, VEC ↥** | ***(91)*** **HUVEC ↓** | **(92) PMVEC ↥** | **(60) HMEC ↥** |

### Monte Carlo calibration improves performance using a single global parameter set

To determine the extent to which validation could be improved without introducing reaction-specific parameters, 100,000 constant-parameter models were generated by Mont Carlo sampling. In each model, a single (*w*, *n*, and *EC_50_*) triplet was assigned globally to the network. Validation accuracy varied substantially across the sampled parameter space (**Fig. 3A**), demonstrating that the qualitative behavior generated by the fixed network topology remained sensitive to the quantitative values assigned to its reactions. Across the 100,000 constant-parameter simulations, mean validation accuracy was 43.6 ± 18.4%.

**Figure 3.**
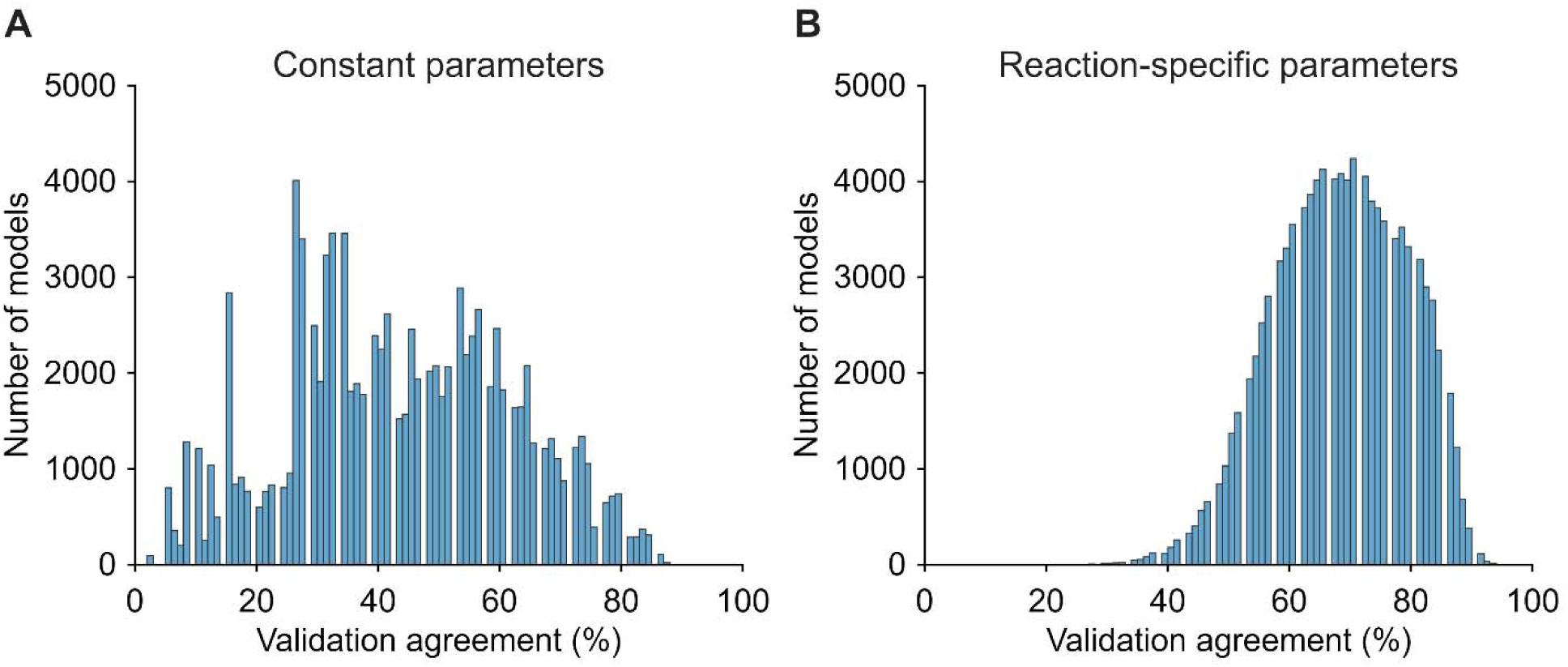
Monte Carlo distributions of validation agreement for globally constant and reaction-specific parameterizations. Validation agreement obtained from 100,000 independently sampled parameter sets for **A.** the constant-parameter model, in which a single *w*, *n*, and *EC_50_*triplet was assigned globally to the network, and **B.** the reaction-specific model, in which independent parameter triplets were assigned to individual reactions. Constant-parameter models yielded a mean validation agreement of 43.6 ± 18.4%, whereas reaction-specific parameterization shifted the distribution toward higher agreement, with a mean of 68.3 ± 10.8%.

The best constant-parameter search reproduced 69 of 79 experimental comparisons, corresponding to 87.34% agreement with the literature validation matrix (**Fig. 4A**) using *w* = 0.772, *n* = 1.599, and *EC_50_* = 0.570. This represented an improvement of seven correctly predicted responses relative to the investigator-selected parameterization. Notably, all defined responses to the stress perturbation were reproduced by the optimized constant model. The remaining ten discrepancies were concentrated within the WSS, hypoxia, and TNFα responses. Under increased WSS, discrepancies remained for ppET1, ET1, pSmad2/3, and proliferation. Under hypoxia, the model did not predict the responses of ET1, pSmad2/3, and apoptosis, while the TNFα condition retained discrepancies in ET1 and proliferation.

**Figure 4.**
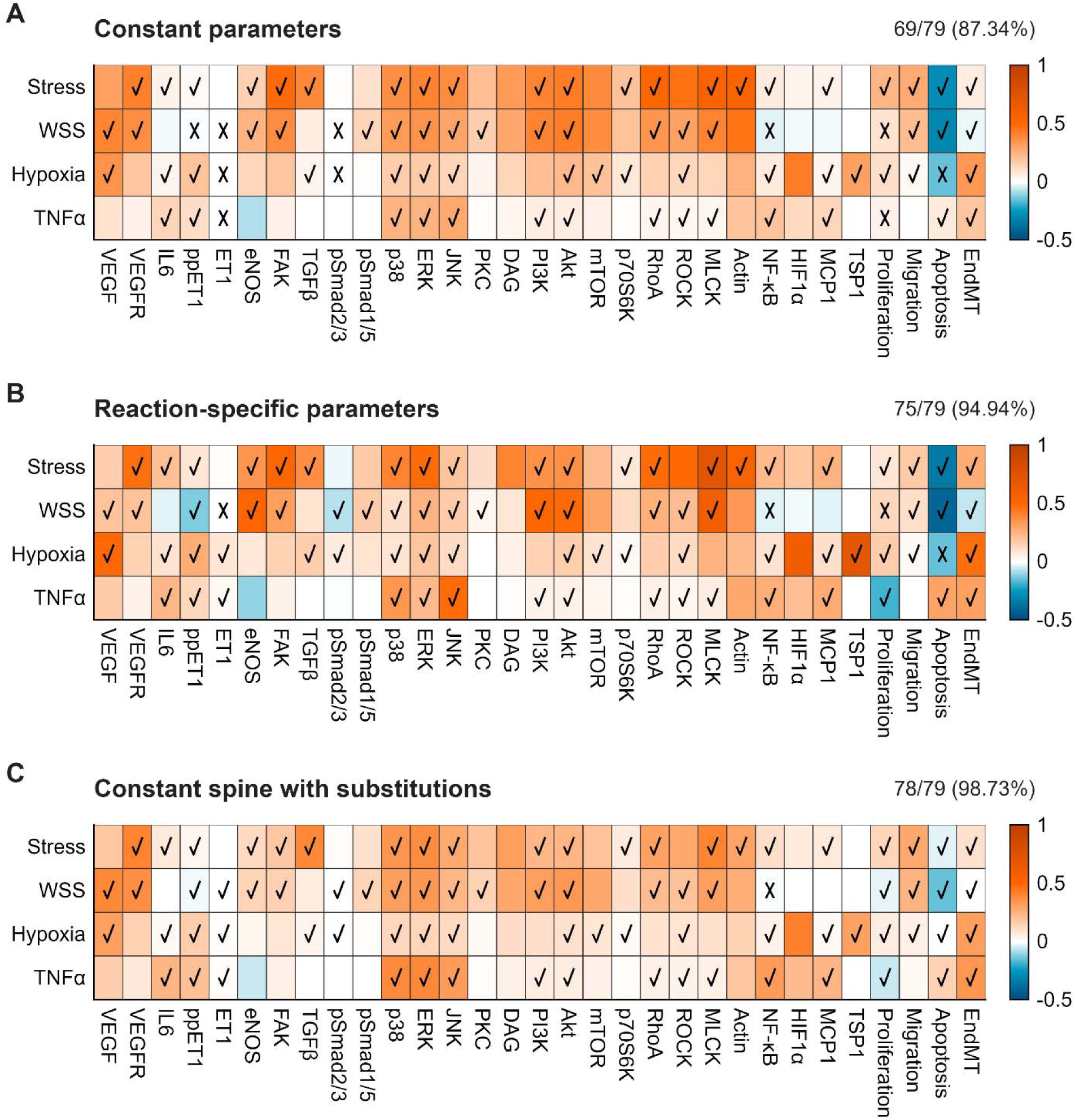
Progressive improvement in endothelial signaling predictions with increasing parameter specificity. Heatmaps compare the highest-performing models obtained using **A.** globally constant parameters, in which the optimized constant-parameter model *w*=0.772, *n*=1.599, *EC_50_*=0.570 reproduced 69 of 79 comparisons (87.34%). **B.** The highest-performing reaction-specific model reproduced 75 of 79 comparisons (94.94%). **C.** Genetic-algorithm optimization of a constant spine with only 18 reaction-specific parameter substitutions reproduced 78 out of 79 validation comparisons (98.73%). Background color denotes the simulated change in normalized activity relative to baseline, while checkmarks and crosses indicate agreement and disagreement, respectively, with the literature-defined directional response.

These results indicate that optimization of a single global reaction parameter set substantially improved the predictive capacity of the fixed network architecture (from 79.5% to 87.34%). Nevertheless, no global parameterization within the sampled space was sufficient to reproduce all literature-defined responses.

### Reaction-specific parameterization further increases agreement with experimental observations

We next tested whether allowing quantitative differences among individual reactions could resolve discrepancies that persisted under the globally constant parameterization. In the perturbed-parameter Monte Carlo analysis, independent values of *w*, *n*, and *EC_50_* were assigned to individual reactions within their prescribed sampling bounds. Across 100,000 parameterizations, the distribution of validation scores shifted toward higher agreement relative to the constant-parameter models (**Fig. 3B**). Whereas the constant-parameter search produced a broad distribution of validation scores, reaction-specific sampling generated a distribution concentrated at substantially higher levels of agreement with the experimental validation matrix. Across the 100,000 reaction-specific parameterizations, mean validation accuracy was 68.3 ± 10.8%.

The highest-performing reaction-specific model reproduced 75 of the 79 directional comparisons, corresponding to 94.94% agreement (**Fig. 4B**). The corresponding reaction-specific parameter values are listed in **Supplemental Table 1**. Reaction-specific parameterization corrected six of the ten discrepancies present in the best constant-parameter model. The remaining disagreements were restricted to three comparisons: ET1 and proliferation responses to increased WSS and the apoptotic response to hypoxia. Responses to stress and TNFα were reproduced completely within the available validation comparisons.

The improvement from 88.46% to 94.94%, despite an identical network topology, indicates that allowing reaction-specific parameter values was necessary to reproduce several experimental responses. Nevertheless, the large number of independently varied parameters raised the question of whether reaction-specific parameters were necessary throughout the network or only within a limited subset of reactions.

### Reaction-specific substitutions achieve near-complete validation of the network

The optimized constant-spine/reaction-specific model achieved 98.73% agreement with the literature validation matrix, correctly reproducing 78/79 defined directional responses (**Fig. 4C**). **Table 2** contains the triplets for the constant spine, as well as the perturbed parameters that improved the accuracy to 98.73%. Near complete agreement required substitution of only 18 reaction parameter triplets from the reaction-specific model, with the remaining reactions retaining their common constant-spine parameterization. These substitutions were sufficient to resolve the four discrepancies that remained in the fully perturbed Monte Carlo solution, including the ET1 and proliferation responses to WSS and the apoptotic response to hypoxia.

**Table 2.** Constant-spine plus reaction-specific parameter values for the optimized endothelial signaling model. The constant-spine parameter triplet (*w* = 0.818, *n* = 1.438, and *EC_50_* = 0.594) was assigned to all reactions except the 18 listed, which used optimized reaction-specific parameter values. For each substituted reaction, the Rpar index, source species, target species, regulatory interaction, and corresponding *w*, *n*, and *EC_50_* values are reported.

| *Rpar index* | *Source* | *Target* | *Regulatory Term* | *w* | *n* | *EC<sub>50</sub>* |
|---|---|---|---|---|---|---|
| **Constant Spine** | - | - | - | 0.818 | 1.438 | 0.594 |
| **Reaction-Specific** |  |  |  |  |  |  |
| **17** | ppET1 | bigET1 | Activation | 0.697 | 2.993 | 0.328 |
| **35** | Integrins | TGFβ | Activation | 0.370 | 2.808 | 0.327 |
| **36** | latent TGFβ | TGFβ | Activation | 0.902 | 2.725 | 0.342 |
| **37** | Stress | latent TGFβ | Activation | 0.896 | 2.497 | 0.451 |
| **38** | latent TGFβ | TGFβ | Activation | 0.639 | 1.489 | 0.452 |
| **39** | TSP1 | TGFβ | Activation | 0.789 | 1.083 | 0.494 |
| **41** | TGFβ | ALK5 | Activation | 0.699 | 2.666 | 0.414 |
| **44** | ALK5 | pSmad2/3 | Activation | 0.379 | 2.489 | 0.564 |
| **69** | TNFαR | ERK | Activation | 0.451 | 2.238 | 0.714 |
| **136** | FAK | PI3K | Activation | 0.517 | 1.255 | 0.432 |
| **138** | Stress | VEGFR | Activation | 0.268 | 2.518 | 0.718 |
| **139** | PI3K | KLF2 | Activation | 0.502 | 1.727 | 0.422 |
| **144** | p38 | p27 | Activation | 0.923 | 1.557 | 0.461 |
| **145** | VEGFR | ERK | Activation | 0.286 | 2.899 | 0.722 |
| **146** | WSS | Notch | Activation | 0.751 | 1.477 | 0.417 |
| **147** | Notch | p27 | Activation | 0.840 | 2.787 | 0.451 |
| **149** | HIF2α | TSP1 | Activation | 0.710 | 1.676 | 0.389 |
| **160** | ALK1 | pSmad1/5 | Activation | 0.952 | 1.433 | 0.533 |

Together, the sequential improvement from 79.49% agreement with the investigator-selected parameterization, to 88.46% following global Monte Carlo calibration, to 96.15% with reaction-specific parameterization, and finally to 98.73% with the final optimized model demonstrates that the majority of endothelial signaling behavior was encoded by the network topology. Near-complete reproduction of the available experimental observations required reaction-specific parameterization of a comparatively small subset of signaling interactions.

### Continuous WSS simulations reveal nonlinear and pathway-dependent endothelial responses

The optimized model was next used to examine endothelial cell responses across a continuous range of normalized WSS. Increasing WSS produced distinct response profiles among the evaluated signaling species and phenotypic mediators (**Fig. 5**). Nitric oxide increased monotonically with WSS and progressively approached a plateau, whereas ET1 exhibited a strongly nonmonotonic response, increasing to a maximum at intermediate WSS before declining at higher levels. The two Smad pathways also responded differently. Phosphorylated Smad2/3 (pSmad2/3) increased modestly at low WSS but progressively decreased as WSS increased, whereas phosphorylated Smad1/5 (pSmad1/5) increased throughout the simulated range. NF-κB similarly exhibited a nonmonotonic response, increasing at low-to-intermediate WSS before decreasing at higher WSS. Thus, although each response arose from the same increase in mechanical input, the optimized signaling network generated markedly different dose-response behaviors across pathways, including monotonic, saturating, and nonmonotonic responses.

**Figure 5.**
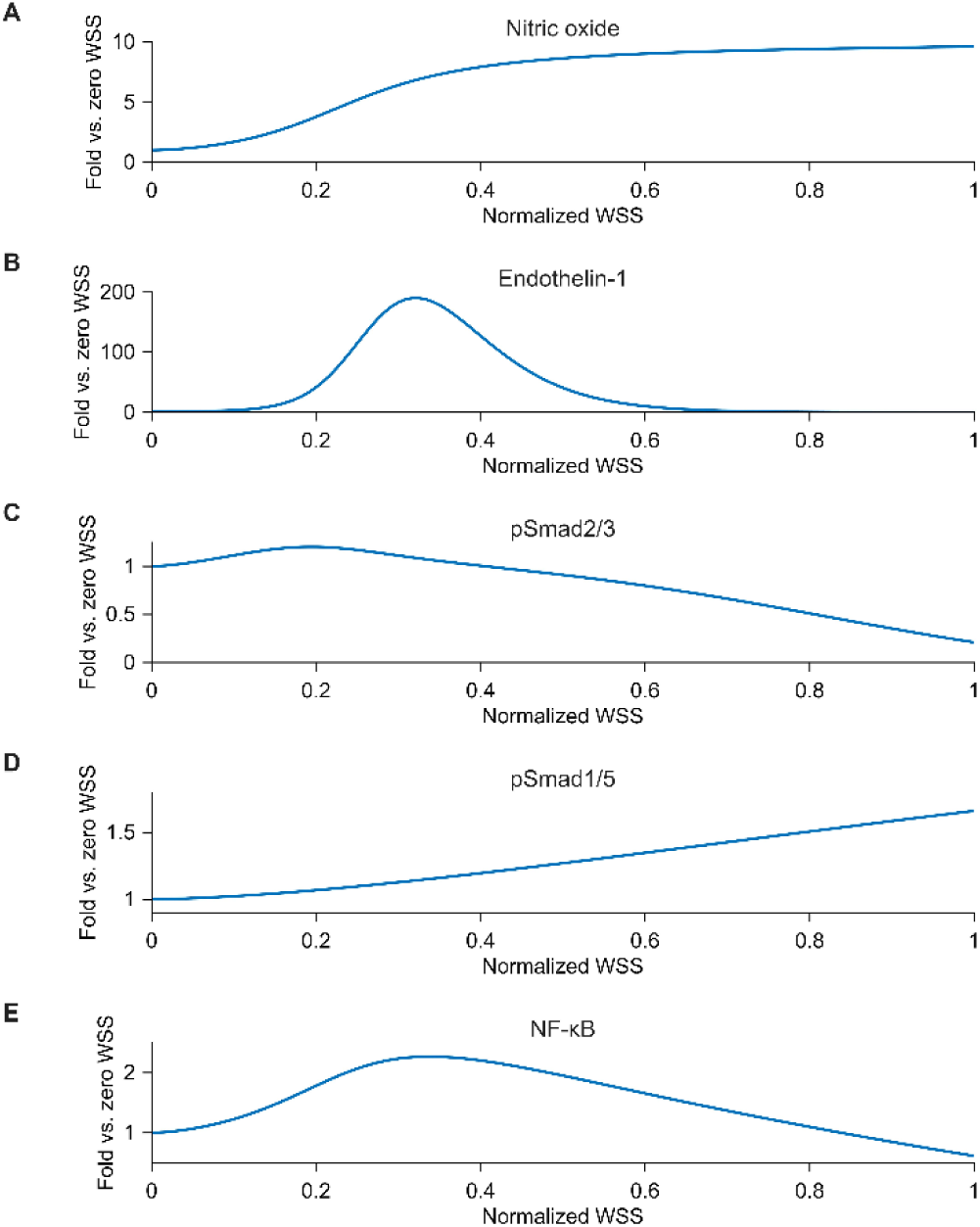
Continuous wall shear stress simulations reveal nonlinear and pathway-dependent endothelial responses. Model-predicted responses to increasing normalized wall shear stress (WSS), expressed as fold change relative to the zero-WSS state, for (A) nitric oxide (NO), (B) Endothelin-1 (ET1), (C) phosphorylated Smad2/3 (pSmad2/3), (D) phosphorylated Smad1/5 (pSmad1/5), and (E) NF-κB. NO increased monotonically and approached a plateau with increasing WSS, whereas ET1 exhibited a strongly nonmonotonic response with a maximum at intermediate WSS. pSmad2/3 increased modestly at low WSS before progressively decreasing, while pSmad1/5 increased throughout the WSS range. NF-κB similarly exhibited a nonmonotonic response, increasing at low-to-intermediate WSS before decreasing at higher WSS. The simulations illustrate that a common mechanical stimulus can generate distinct dose-response behaviors across endothelial signaling pathways.

## Discussion

Computational models of vascular mechanics and mechanobiology continue to provide considerable insight into a host of adaptations to sustained changes in pressure and flow as well as diverse cases of disease progression (93) and, recently, even natural postnatal development (94). Most of these models are phenomenological, based on constrained mixture theory (95), and thus depend on constituent-specific constitutive relations for mass production (matrix deposition, cell proliferation), mass removal (matrix degradation, cell death), and nonlinear material behaviors. Despite the valuable knowledge gained, these models do not provide mechanistic understanding. For this reason, cell signaling models have been developed and coupled to tissue-level phenomenological models to enable simulations of changing geometry, composition, and properties to be linked to cell-level signaling (96, 97). Although the phenomenological models capture certain mechanobiological effects of endothelial cells, there had previously been a lack of an endothelial cell signaling model sufficient for coupling with tissue-level models. In this paper, we present an endothelial cell signaling model built from the archival literature. As seen in **Fig. 1**, the network contains many of the key signaling pathways, including RhoA/ROCK, MAPK, Smad, PI3K-Akt-mTOR, and JAK-STAT, among others.

The good agreement between network and data for the globally constant-parameter (*w*, *n*, *EC_50_*) model suggests that network topology was appropriate. Namely, the optimized constant-parameter model reproduced 69 of 79 literature-defined responses, including all responses to intramural stress and most responses to shear, hypoxia, and TNFα. That much of the qualitative behavior of the endothelial cell could be captured by network connectivity and regulatory direction alone, without requiring unique parameterization of individual reactions, is consistent with many prior signaling models in which a biologically meaningful network can emerge using broadly shared or default reaction parameters, despite limited availability of reaction-specific kinetic data (3, 9, 11).

Importantly, we additionally used Monte Carlo simulations and a genetic algorithm optimization to determine a limited set of species-specific parameters that increased the fidelity of the model. Prior literature-derived logic-based signaling models have demonstrated substantial qualitative predictive accuracy, including a brain endothelial model that reproduced 73% of independent experimental relationships (5) and a pulmonary arterial fibroblast model that reproduced 80% of independent validation observations without parameter optimization (4). In contrast to these prior approaches, we explicitly tested the extent to which predictive agreement was governed by network topology versus reaction-specific parameterization. Consistent with the aforementioned conclusion that network topology was sufficient, only 18 of the 123 parameters needed to be species-specific to increase the agreement with data to 98.73%, which is unprecedented in similar prior models, including our own (3, 5, 10, 11). Notably, several of the reaction-specific substitutions occurred within pathways central to endothelial mechanotransduction (including VEGFR-to-RhoA, VEGFR-to-PI3K, RhoA-to-FAK, FAK-to-PI3K, and WSS-to-Notch). Thus, the requirement for reaction-specific parameterization at these interactions is consistent with their roles as important points of integration between mechanical stimuli and downstream cell signaling. More broadly, maintaining a common parameter set for the majority of the network while allowing quantitative specialization of a limited subset of reactions reduces the number of independently specified parameters, an important consideration in large biology models in which individual parameter values are otherwise poorly identified (98). The reaction-specific values determined here should therefore be interpreted as effective parameters that improve network-level predictive behavior rather than uniquely identified constants. In summary, our findings suggest that network topology establishes much of the qualitative behavior of the endothelial signaling network; quantitative specialization of selected reactions then adds specificity that cannot be captured using globally shared parameters alone.

Due to the limited number of changes compared to a fully reaction-specific network, the resulting parameterization may be interpretable. For example, the Hill coefficient governs the steepness of the activation function, with lower values producing gradual responses and higher values producing switch-like behavior. The two reactions with the highest values were ppET1 => bigET1 (*n* = 2.993) VEGFR => ERK (*n* = 2.899). These higher values may reflect a more switch-like behavior and therefore better capture the kinetics compared to the lower values of from the constant-spine parameterization. In turn, the weighting parameter provides insight into the relative importance of individual reactions. Stress => VEGFR received the lowest weight (*w* = 0.286), suggesting that this reaction contributes relatively little to the model’s behavior under the fitted conditions. Inspection of the network topology reveals alternative pathways by which mechanical stimuli regulate BBEFR and downstream signaling, potentially indicating some degree of functional redundancy. Alternatively, this result may motivate closer examination of the network structure to determine whether these direct interactions are truly a distinct signaling mechanism or if they simply overlap with indirect pathways. More generally, extreme reductions in may identify reactions whose removal from the network could be considered as candidate topological simplifications.

The continuous WSS simulations demonstrate the ability of the optimized network to generate pathway-specific responses beyond the discrete perturbations used for model validation. Several of these predictions were consistent with experimentally reported responses to different magnitudes of shear stress. Monotonic increases of NO are consistent with the well-established activation of endothelial NO signaling by increased WSS (35). Similarly, a greater Smad2/3 response at low WSS is consistent with Smad2/3 localization and target-gene expression being maximal under low shear and suppressed at higher physiological shear (50). The predicted decrease in NF-κB signaling at higher WSS is consistent with experimental findings that prolonged low shear produces greater NF-κB activation than high laminar shear (99). Not all predictions agreed with available data, however. The model predicted a continued increase in pSmad1/5 with WSS, whereas recent studies have shown that high shear can suppress Smad1/5 signaling (100). This regulatory interaction is not represented in the present network and thus provides one example of how comparison of model-generated dose-response predictions with experimental data can identify signaling mechanisms for future model refinement. Of note, topological changes that produce this behavior can be hypothesized and tested within the model; previous analyses have focused on modifying the network structure to improve validation accuracy (101) and this is likely to be complementary to the focus on parameterization herein.

Notwithstanding advances achieved, several limitations should be considered. The model provides a simplified representation of endothelial biology, with signaling processes represented by lumped nodes and phenomenological reaction equations rather than detailed biochemical kinetics. In addition, model agreement was evaluated primarily by the direction of experimental responses (e.g., up or down regulation) rather than their magnitude or temporal dynamics. Such qualitative assessments can prove especially challenging when nonmonotonic responses exist, as the directional observation may depend on initial conditions or magnitude of perturbation. For species exhibiting these types of behaviors, validation should be extended to consider the appropriate context. Note that we also focused on literature related to laminar shear stress, although distinct behaviors are expected to emerge under oscillatory or disturbed flow. Finally, the validation matrix was assembled from a heterogeneous literature spanning different endothelial cell types, species, experimental conditions, and durations of signaling, noting that similarly diverse literature also informed construction of the signaling network. Thus, the reported agreement should be interpreted primarily as consistency with the available literature rather than fully independent prospective validation. Future directions should include further optimization using data for other experimental conditions, adding crosstalk parameters, and accelerating learning relationships between species and reactions using neural networks.

In conclusion, we presented a novel 70-species intracellular signaling model for the endothelium and showed, using Monte Carlo simulations and methods of optimization, that most of the 123 parameter triplets (*w*, *n*, *EC_50_*) can be assigned the same values while only a fraction needs to be assigned species-specific values. These findings suggest that the majority of qualitative endothelial signaling behavior is encoded by network topology, with quantitative specialization required for only a limited subset of signaling interactions. The resulting model predicted distinct nonlinear responses of key endothelial pathways to continuously varying WSS, several of which were consistent with experimentally reported shear-dependent responses. Together, this framework provides a computationally efficient representation of endothelial signaling that can be extended to additional experimental conditions and ultimately coupled with tissue-level models of vascular adaptation and disease.

## Methods

### Model structure: signaling networks as directed graphs

The intracellular endothelial cell signaling network was represented as a directed graph in which nodes (species) were connected by edges (reactions) with defined directionality, namely activation versus inhibition. Nodes represented extracellular and mechanical inputs, membrane-bound receptors, intracellular signaling intermediates, transcription factors, gene products, and phenotypic outputs. Network topology was assembled from prior literature and endothelial cell-specific signaling knowledge. Importantly, the literature used for network construction was independent from the literature used for validation, as shown in **Table 1**.

The directed graph was converted into a continuous logic-based ordinary differential equation (ODE) model using the normalized-Hill framework implemented in NetFlux (9), consistent with our previous vascular signaling networks (3, 11). Each node *i* was assigned a normalized activity level *Y_i_* ∈[0,1] and its evolution was governed by

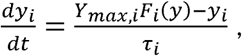

Where *Y_max,i_* ∈ [0,1], 0,1 is the maximum activity of species *i*, τ_i_ is a characteristic time constant, and F_i_(y) is a composite regulatory input function determined by upstream activating and inhibitory reactions affecting species *i*. The graph topology defines which species regulates each node, while the reaction parameters define the strength and shape of those regulatory effects. Each regulatory reaction was characterized by a parameter triplet,

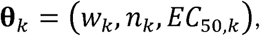

Where *W_k_* is the maximum reaction weight, n_k_ is the Hill coefficient, and EC_50,k_ is the upstream activity producing half of the maximal edge response. For an upstream species with normalized activity *X*, the activating reaction was defined by

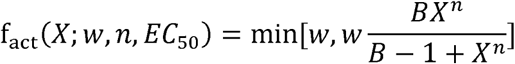

where

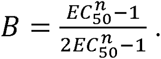

The corresponding inhibitory reaction was defined by

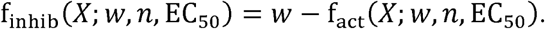

When multiple upstream reactions regulated the same target, their weighted reaction outputs were combined using continuous analogs of Boolean logic. For reaction outputs joined by an AND relationship, the implemented operator was

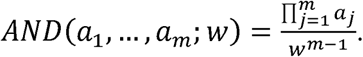

If a and b denote the outputs of two activating or inhibitory Hill functions, the OR operator was

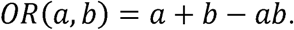

### Ordinary differential equation simulation and experimental validation

All candidate parameterizations were evaluated using common baseline and perturbation conditions. Model inputs were represented as dimensionless activities on a normalized scale from 0 to 1. The intramural Stress, Wall Shear Stress (WSS), Hypoxia, Caveolin-1, and TNF-alpha input states were initialized to 0.2, with stress overridden to 0.5 to establish basal TGF-beta activity. The exogenous angiotension II (AngII) input was inactive, with its input weight fixed to zero, while AngII’s downstream receptors remained active parts of the model’s architecture. The model was first simulated under a baseline initial condition. Stress, WSS, Hypoxia, and TNF-alpha were then tested independently by adding 0.5 to the corresponding input state in the final baseline solution and integrating the perturbed model over additional times. Evaluation of one parameter set therefore required five ODE integrations: one baseline simulation and four perturbation simulations.

For perturbation *p* and species *j*, the predicted change was calculated from the difference between perturbed and baseline states, computed at the final timepoint t*_f_*, via

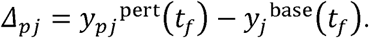

The continuous change was converted into a qualitative directional prediction and compared with the literature-derived validation matrix. The presented validation matrix contained 79 defined directional comparisons. The acceptable range for no change was an interval of ±0.02 around a change of 0.

### Monte Carlo Optimization

Monte Carlo analysis was used to calibrate and evaluate the fixed endothelial signaling network. It did not infer network topology, as topology was set by literature-derived relationships of species and reactions, as previously mentioned. In this instance, Monte Carlo sampling refers to independent random searches over prespecified parameter bounds. Monte Carlo searches used 100,000 parameter sets. To keep the normalized Hill function real-valued when *n* was noninteger, the upper sampling limit for EC_50_ was restricted to

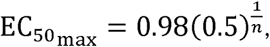

which was chosen as a practical implementation of the inequality EC^n^_50_ < 0.5, which ensures B – 1 > 0. The Monte Carlo simulations were performed in MATLAB R2025a (MathWorks, Inc., Natick, MA, USA) using the Parallel Computing Toolbox.

1. *Constant Parameter Model:* The constant-parameter search sought the best performance achievable using a single parameter triplet for all reactions. In each of 100,000 runs, one global parameter triplet (*W,n,EC_50_*) was sampled and assigned to every reaction in the network, except that the exogenous AngII input weight remained zero. The sampled ranges were

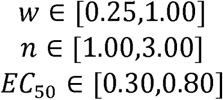

subject to the n-dependent EC_50_ limit above. Every sampled triplet was evaluated with the common baseline and four-condition validation procedure.

2. *Perturbed Parameter Model:* The edge-perturbed search tested whether allowing differences in parameter values among reactions improved the ability of the same network topology to reproduce the literature based endothelial signaling responses. A separate triplet (*W,n,EC_50_*) was sampled independently for every reaction in each of 100,000 runs. For reactions without manually specified bounds, the default ranges were the same as the constant parameter model.

3. *Constant-Spine with reaction-specific perturbations:* Although the edge-perturbed search provided more flexibility, a fully reaction-specific model assigns three independently sampled parameters to every reaction. Such an approach risks overparameterizing the network and also limits the ability to mechanistically interpret the most important parameter changes. A constant-spine with reaction-specific analysis was thus developed to retain the simplicity of the constant parameter model while identifying the smallest set of reactions for which edge-specific values were needed to achieve the highest available validation accuracy. This stage selected among parameter values generated by the preceding searches; it did not sample new continuous parameter values.

All constant-parameter models that tied for the highest validation score were retained as candidate spines. Reaction-specific perturbations were selected from the 50 saved highest-ranked edge-perturbed models. Twenty constant spines were paired with 10 retained reaction-specific models, producing 200 candidate combinations.

For each constant-spine/reaction-specific model pair, a binary variable *z_k_* ∈ [0,1] was assigned to every reaction eligible for substitution. The parameter triplet used for reaction *k* was

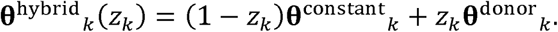

Thus, *z_k_* =0 retained the constant-spine triplet and Z_k_ =1 substituted the complete (*W_k_,n_k_,EC_50k_*) triplet from the paired reaction-specific models. All substituted reactions within one candidate model came from the same perturbed model; parameters from different perturbed runs were not mixed edge by edge. The present network contained 123 reaction positions eligible for this binary selection.

Reaction-specific substitutions were then removed individually when their removal did not reduce validation accuracy, with this procedure repeated up to two passes. Screened pairs were ranked first by validation matches and then by the number of reaction-specific substitutions, with the five highest-ranked pairs advanced to the final optimization. Each finalist pair was evaluated with a binary genetic algorithm using five independent random seeds, producing 25 finalist searches. The population contained 250 candidate substitution patterns and was evolved for 200 generations. The initial population included the all-constant substitution pattern, the all-reaction-specific substitution pattern, the substitution pattern obtained during the initial screening procedure, and randomly generated models containing relatively few reaction-specific substitutions, with reaction-specific inclusion probabilities of 0.02, 0.05, 0.10, 0.20, or 0.35. The five highest-ranked solutions were retained at each generation. Parent solutions were selected by comparing three candidates and retaining the highest-ranked. Offspring inherited each binary value with equal probability from either parent, and each value had a 3% probability of mutation.

The genetic algorithm maximized

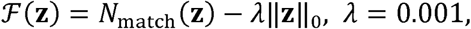

Where ‖**z**‖_0_ = Σ_*k*_*z*_*k*_ is the number of reaction-specific substitutions. The complexity penalty was smaller than the value of one additional validation match across the full range of possible model sizes. As a result, validation accuracy was always optimized first, while model size was used to distinguish models having the same accuracy. Duplicate substitution patterns were cached to avoid repeated simulations. After the final generation, the highest-ranked mask was further pruned for up to two passes.

Final models were ranked first by maximum validation accuracy, then by the minimum number of reaction-specific substitutions. Among equally accurate final models, we recorded how often each reaction was replaced and the range of its parameter values to identify reactions that were consistently selected versus those that varied among models.

## Acknowledgments

This research was supported, in part, by grants from the National Institutes of Health under grants R01 HL169147 and P01 HL169168 (J.D.H.).

