## Supplemental Data for "From Network Topology to Parameterization: An Optimized Model of Endothelial Mechanobiology"

Pete H. Gueldner^1^, Linda Irons^1,+^, Jeremy L. Warren^1^, David S. Li^1,*^, Jay D. Humphrey^1,2^

*^1^Department of Biomedical Engineering, Yale University, New Haven, CT, USA*

*^2^Vascular Biology and Therapeutics Program, Yale School of Medicine, New Haven, CT, USA*

*^+^Current Address: AstraZeneca, Gaithersburg, MD, USA*

*^*^Current Address: Department of Biomechanics, University of Nebraska at Omaha, Omaha, NE, USA*

Address correspondence to:

Pete H. Gueldner, Ph.D.

Department of Biomedical Engineering

Yale University

**Supplementary Table 1. Reaction-specific parameter values for the selected highest-performing model.** The network contained 123 reaction rules represented by 152 active parameterized regulatory edges and logic-gate terms. Values correspond to Monte Carlo run 62672, selected as the first of two parameter sets tied for the highest validation score. For each parameterized term, *w* denotes reaction weight, *n* denotes the Hill coefficient, and *EC_50_* denotes the normalized upstream activity producing a half-maximal response.

|  | *Rpar index* | *Source* | *Target* | *Regulatory term* | *w* | *n* | *EC_50_* | *References* |
| --- | --- | --- | --- | --- | --- | --- | --- | --- |
|  | 8 | eNOS | NO | Activation and AND gate | 0.482 | 2.207 | 0.369 | (1) |
|  | 9 | ROS | NO | Inhibition | 0.245 | 2.089 | 0.305 | (1, 2) |
|  | 10 | Stress | IonChannels | Activation | 0.409 | 1.359 | 0.482 | (3) |
|  | 11 | Wss | IonChannels | Activation | 0.775 | 2.809 | 0.452 | (4) |
|  | 12 | IonChannels | Ca | Activation | 0.805 | 1.397 | 0.517 | (4, 5) |
|  | 13 | Ca | CaM | Activation | 0.746 | 1.605 | 0.366 | (6, 7) |
|  | 14 | CaM | eNOS | Activation and AND gate | 0.467 | 1.057 | 0.446 | (8, 9) |
|  | 15 | Cav1 | eNOS | Inhibition | 0.710 | 2.396 | 0.409 | (8, 9) |
|  | 16 | TNFaR | eNOS | Inhibition | 0.885 | 2.151 | 0.693 | (8, 9) |
|  | 17 | ppET1 | bigET1 | Activation | 0.553 | 2.843 | 0.433 | (10) |
|  | 18 | bigET1 | ECE | Activation | 0.910 | 1.877 | 0.326 | (10) |
|  | 19 | ECE | ET1 | Activation | 0.254 | 1.687 | 0.565 | (10) |
|  | 20 | ET1 | ETBR | Activation | 0.819 | 1.571 | 0.509 | (11) |
|  | 21 | ETBR | PI3K | Activation | 0.906 | 2.588 | 0.432 | (12) |
| 22 | | Wss | ETBR | Activation | 0.785 | 1.782 | 0.583 | (10) |
|  | 23 | Hypoxia | HIF1a | Activation | 0.891 | 2.260 | 0.450 | (13-15) |
|  | 24 | HIF1a | ppET1driver | Activation | 0.522 | 1.829 | 0.564 | (14) |
|  | 25 | pSmad23 | ppET1driver | Activation | 0.975 | 1.058 | 0.349 | (16, 17) |
|  | 26 | NFkB | ppET1driver | Activation | 0.292 | 1.785 | 0.400 | (18, 19) |
|  | 27 | AT1R | ppET1driver | Activation | 0.717 | 2.630 | 0.746 | (20) |
|  | 28 | ppET1driver | ppET1 | Activation and AND gate | 0.335 | 1.073 | 0.374 | (10) |
|  | 29 | NO | ppET1 | Inhibition | 0.685 | 2.877 | 0.552 | (10) |
|  | 30 | Akt | eNOS | Activation and AND gate | 0.847 | 1.560 | 0.333 | (9, 21-24) |
|  | 31 | RhoA | eNOS | Inhibition | 0.995 | 1.884 | 0.443 | (9, 21-24) |
|  | 32 | Cav1 | eNOS | Inhibition | 0.282 | 2.338 | 0.383 | (9, 21-24) |
|  | 33 | TNFaR | eNOS | Inhibition | 0.830 | 2.448 | 0.562 | (9, 21-24) |
|  | 34 | Stress | Integrins | Activation | 0.668 | 2.279 | 0.422 | (25) |
|  | 35 | Integrins | TGFb | Activation and AND gate | 0.561 | 1.950 | 0.341 | (26) |
|  | 36 | latentTGFb | TGFb | Activation | 0.937 | 1.265 | 0.407 | (27) |
|  | 37 | Stress | latentTGFb | Activation | 0.665 | 1.662 | 0.521 | (28, 29) |
|  | 38 | latentTGFb | TGFb | Activation and AND gate | 0.459 | 2.634 | 0.309 | (27) |
|  | 39 | TSP1 | TGFb | Activation | 0.513 | 1.465 | 0.318 | (27) |
|  | 40 | TGFb | ALK1 | Activation | 0.500 | 1.358 | 0.492 | (30, 31) |
|  | 41 | TGFb | ALK5 | Activation | 0.556 | 1.193 | 0.447 | (30, 31) |
|  | 42 | ALK1 | pSmad15 | Activation and AND gate | 0.936 | 2.376 | 0.413 | (30, 31) |
|  | 43 | Wss | pSmad15 | Activation | 0.957 | 1.922 | 0.677 | (30, 31) |
|  | 44 | ALK5 | pSmad23 | Activation and AND gate | 0.858 | 2.578 | 0.309 | (31-34) |
|  | 45 | Klf2 | pSmad23 | Inhibition | 0.898 | 1.315 | 0.520 | (31-34) |
|  | 46 | VEGFR | pSmad23 | Inhibition | 0.955 | 1.505 | 0.358 | (31-34) |
|  | 47 | pSmad23 | PAI1 | Activation | 0.930 | 1.378 | 0.372 | (31, 35) |
|  | 48 | pSmad23 | p38 | Activation | 0.907 | 2.644 | 0.587 | (36) |
|  | 49 | pSmad15 | Id1 | Activation | 0.291 | 1.139 | 0.444 | (30) |
|  | 50 | Id1 | Migration | Activation and AND gate | 0.721 | 2.308 | 0.486 | (30) |
|  | 51 | PAI1 | Migration | Inhibition | 0.944 | 1.743 | 0.632 | (30) |
|  | 52 | pSmad15 | Proliferation | Activation and AND gate | 0.483 | 2.928 | 0.337 | (30) |
|  | 53 | TGFb | Proliferation | Activation | 0.292 | 2.182 | 0.553 | (30) |
|  | 54 | p27 | Proliferation | Inhibition | 0.294 | 2.488 | 0.433 | (30) |
|  | 55 | pSmad23 | EndMT | Activation | 0.923 | 1.503 | 0.532 | (37, 38) |
|  | 56 | PI3K | Akt | Activation | 0.959 | 1.108 | 0.414 | (39) |
|  | 57 | Akt | mTOR | Activation | 0.560 | 1.433 | 0.579 | (39) |
|  | 58 | mTOR | mTORC1 | Activation | 0.563 | 1.709 | 0.418 | (39) |
|  | 59 | mTOR | mTORC2 | Activation | 0.895 | 2.318 | 0.621 | (39) |
|  | 60 | mTORC2 | Akt | Activation | 0.545 | 2.694 | 0.746 | (39) |
|  | 61 | mTORC1 | p70S6K | Activation | 0.447 | 1.919 | 0.441 | (40) |
|  | 62 | p70S6K | Proliferation | Activation and AND gate | 0.252 | 1.063 | 0.449 | (40) |
|  | 63 | p27 | Proliferation | Inhibition | 0.454 | 2.643 | 0.485 | (40) |
|  | 64 | Akt | NFkB | Activation and AND gate | 0.244 | 2.780 | 0.605 | (41, 42) |
|  | 65 | Klf2 | NFkB | Inhibition | 0.622 | 1.603 | 0.452 | (41, 42) |
|  | 66 | TNFa | TNFaR | Activation | 0.796 | 1.210 | 0.437 | (43) |
|  | 67 | TNFaR | JNK | Activation | 0.995 | 2.258 | 0.537 | (44, 45) |
|  | 68 | TNFaR | p38 | Activation | 0.774 | 1.428 | 0.566 | (44, 45) |
|  | 69 | TNFaR | ERK | Activation | 0.676 | 2.166 | 0.675 | (46) |
|  | 70 | TNFaR | p27 | Activation | 0.852 | 2.175 | 0.437 | (47) |
|  | 71 | TNFaR | NFkB | Activation and AND gate | 0.931 | 1.458 | 0.565 | (42, 43) |
|  | 72 | Klf2 | NFkB | Inhibition | 0.642 | 1.400 | 0.483 | (42, 43) |
|  | 73 | IL6 | IL6R | Activation | 0.578 | 2.430 | 0.413 | (48) |
|  | 74 | NFkB | IL6 | Activation | 0.656 | 1.051 | 0.321 | (43) |
|  | 75 | IL6R | JAK | Activation and AND gate | 0.988 | 2.414 | 0.549 | (48-50) |
|  | 76 | SOCS | JAK | Inhibition | 0.514 | 2.025 | 0.682 | (48-50) |
|  | 77 | JAK | PI3K | Activation | 0.822 | 2.984 | 0.601 | (48, 49) |
|  | 78 | JAK | STAT3 | Activation | 0.813 | 1.749 | 0.471 | (48, 49) |
|  | 79 | STAT3 | SOCS | Activation | 0.744 | 1.213 | 0.486 | (48, 51) |
|  | 80 | JAK | ERK | Activation | 0.933 | 1.976 | 0.616 | (48, 49) |
|  | 81 | STAT3 | MCP1 | Activation and AND gate | 0.413 | 2.859 | 0.736 | (49) |
|  | 82 | PI3K | MCP1 | Activation | 0.751 | 1.294 | 0.500 | (49) |
|  | 83 | NFkB | EndMT | Activation | 0.454 | 2.014 | 0.325 | (52) |
|  | 84 | NFkB | HIF1a | Activation | 0.628 | 2.118 | 0.428 | (53, 54) |
|  | 85 | NFkB | MCP1 | Activation | 0.572 | 1.832 | 0.311 | (55-57) |
|  | 86 | HIF1a | VEGF | Activation | 0.691 | 1.732 | 0.492 | (37, 58) |
|  | 87 | Klf2 | VEGF | Activation | 0.340 | 2.535 | 0.693 | (59) |
|  | 88 | VEGF | VEGFR | Activation | 0.397 | 1.506 | 0.535 | (60, 61) |
|  | 89 | VEGFR | RhoA | Activation | 0.644 | 2.138 | 0.33 | (62) |
|  | 90 | VEGFR | PI3K | Activation | 0.642 | 1.051 | 0.451 | (39) |
|  | 91 | VEGFR | PLCγ | Activation | 0.875 | 2.465 | 0.596 | (1, 60, 63, 64) |
|  | 92 | VEGFR | p38 | Activation | 0.849 | 2.945 | 0.681 | (65, 66) |
|  | 93 | VEGFR | FAK | Activation | 0.591 | 1.343 | 0.493 | (67) |
|  | 94 | PLCγ | DAG | Activation | 0.750 | 2.220 | 0.552 | (1, 68) |
|  | 95 | DAG | PKC | Activation and AND gate | 0.75 | 1.397 | 0.358 | (1, 68) |
|  | 96 | Ca | PKC | Activation | 0.575 | 2.252 | 0.625 | (1, 68) |
|  | 97 | PKC | NFkB | Activation and AND gate | 0.322 | 2.315 | 0.321 | (24, 42, 69, 70) |
|  | 98 | Klf2 | NFkB | Inhibition | 0.924 | 1.497 | 0.476 | (24, 42, 69, 70) |
|  | 99 | PKC | eNOS | Activation and AND gate | 0.320 | 1.028 | 0.432 | (9, 23, 68) |
|  | 100 | RhoA | eNOS | Inhibition | 0.633 | 2.998 | 0.501 | (9, 23, 68) |
|  | 101 | Cav1 | eNOS | Inhibition | 0.360 | 2.216 | 0.539 | (9, 23, 68) |
|  | 102 | TNFaR | eNOS | Inhibition | 0.969 | 2.531 | 0.483 | (9, 23, 68) |
|  | 103 | PKC | ERK | Activation | 0.821 | 1.954 | 0.549 | (71) |
|  | 104 | PLCγ | Ca | Activation | 0.390 | 1.137 | 0.418 | (64) |
|  | 105 | CaM | MLCK | Activation | 0.693 | 2.505 | 0.345 | (6, 7) |
|  | 106 | RhoA | ROCK | Activation | 0.864 | 2.572 | 0.649 | (72) |
|  | 107 | RhoA | FAK | Activation | 0.733 | 2.286 | 0.643 | (73) |
|  | 108 | ROCK | Actin | Activation | 0.486 | 1.357 | 0.402 | (72) |
|  | 109 | ROCK | MLCK | Activation | 0.986 | 1.439 | 0.316 | (72) |
|  | 110 | MLCK | Myosin | Activation | 0.534 | 1.822 | 0.473 | (6, 74) |
|  | 111 | Actin | Migration | Activation and AND gate | 0.360 | 2.225 | 0.602 | (31) |
|  | 112 | Myosin | Migration | Activation | 0.265 | 2.784 | 0.386 | 28 |
|  | 113 | PAI1 | Migration | Inhibition | 0.461 | 2.633 | 0.459 | 28 |
|  | 114 | p38 | Actin | Activation | 0.630 | 1.276 | 0.395 | (65, 66) |
|  | 115 | ERK | Proliferation | Activation and AND gate | 0.853 | 1.950 | 0.313 | (46, 75) |
|  | 116 | p27 | Proliferation | Inhibition | 0.760 | 2.359 | 0.567 | (46, 75) |
|  | 117 | JNK | BAD | Activation and AND gate | 0.581 | 2.197 | 0.511 | (76-79) |
|  | 118 | Akt | BAD | Inhibition | 0.813 | 1.851 | 0.305 | (76-79) |
|  | 119 | JNK | foxo | Activation and AND gate | 0.626 | 1.131 | 0.465 | (6, 76, 77) |
|  | 120 | Akt | foxo | Inhibition | 0.927 | 2.685 | 0.414 | (6, 76, 77) |
|  | 121 | BasalDeath | BAD | Activation and AND gate | 0.602 | 1.694 | 0.592 | (78-80) |
|  | 122 | Akt | BAD | Inhibition | 0.296 | 2.281 | 0.593 | (78-80) |
|  | 123 | BasalDeath | foxo | Activation and AND gate | 0.455 | 1.475 | 0.361 | (77, 80) |
|  | 124 | Akt | foxo | Inhibition | 0.716 | 1.686 | 0.358 | (77, 80) |
|  | 125 | Wss | Integrins | Activation | 0.890 | 2.799 | 0.349 | (81-83) |
|  | 126 | Integrins | RhoA | Activation and AND gate | 0.761 | 2.955 | 0.679 | (84) |
|  | 127 | Stress | RhoA | Activation | 0.652 | 2.530 | 0.691 | (84) |
|  | 128 | Integrins | FAK | Activation and AND gate | 0.738 | 1.825 | 0.574 | (85, 86) |
|  | 129 | Stress | FAK | Activation | 0.448 | 2.859 | 0.355 | (85, 86) |
|  | 130 | FAK | Cdc42 | Activation | 0.863 | 2.792 | 0.345 | (87) |
|  | 131 | Cdc42 | JNK | Activation | 0.379 | 1.757 | 0.430 | (88) |
|  | 132 | Cdc42 | Actin | Activation | 0.395 | 2.112 | 0.598 | (89) |
|  | 133 | BAD | Apoptosis | Activation | 0.610 | 2.596 | 0.427 | (78, 79) |
|  | 134 | foxo | Apoptosis | Activation | 0.955 | 2.239 | 0.364 | (77) |
|  | 135 | HIF1a | EndMT | Activation | 0.686 | 1.316 | 0.472 | (90) |
|  | 136 | FAK | PI3K | Activation | 0.261 | 2.491 | 0.378 | (91) |
|  | 137 | Wss | VEGFR | Activation | 0.296 | 2.406 | 0.465 | (92) |
|  | 138 | Stress | VEGFR | Activation | 0.692 | 1.823 | 0.651 | (93) |
|  | 139 | PI3K | Klf2 | Activation | 0.552 | 2.138 | 0.700 | (42, 94) |
|  | 140 | Klf2 | eNOS | Activation and AND gate | 0.264 | 1.969 | 0.680 | (9, 23, 24, 42) |
|  | 141 | RhoA | eNOS | Inhibition | 0.409 | 2.417 | 0.449 | (9, 23, 24, 42) |
|  | 142 | Cav1 | eNOS | Inhibition | 0.260 | 1.871 | 0.480 | (9, 23, 24, 42) |
|  | 143 | TNFaR | eNOS | Inhibition | 0.883 | 2.534 | 0.706 | (9, 23, 24, 42) |
|  | 144 | p38 | p27 | Activation | 0.773 | 1.04 | 0.355 | (88, 95) |
|  | 145 | VEGFR | ERK | Activation | 0.882 | 1.70 | 0.394 | (88) |
|  | 146 | Wss | Notch | Activation | 0.495 | 1.677 | 0.643 | (75) |
|  | 147 | Notch | p27 | Activation | 0.638 | 2.363 | 0.445 | (75) |
|  | 148 | Hypoxia | HIF2a | Activation | 0.873 | 2.407 | 0.509 | (96) |
|  | 149 | HIF2a | TSP1 | Activation | 0.858 | 1.873 | 0.302 | (96) |
|  | 150 | Wss | Klf2 | Activation | 0.982 | 2.372 | 0.563 | (94, 97) |
|  | 151 | AngIIin | AT1R | Activation | 0.279 | 1.394 | 0.531 | (98) |
|  | 152 | AngIIin | AT2R | Activation | 0.700 | 2.497 | 0.422 | (99) |
|  | 153 | AT1R | NOX2 | Activation | 0.912 | 2.870 | 0.639 | (100) |
|  | 154 | NOX2 | ROS | Activation | 0.316 | 1.849 | 0.674 | (100) |
|  | 155 | ROS | NFkB | Activation | 0.275 | 2.213 | 0.691 | (101) |
|  | 156 | AT1R | RhoA | Activation | 0.621 | 1.463 | 0.406 | (98) |
|  | 157 | AT2R | eNOS | Activation | 0.967 | 1.865 | 0.490 | (99) |
|  | 159 | BMP9_10 | ALK1 | Activation | 0.276 | 1.750 | 0.353 | (32) |
|  | 160 | ALK1 | pSmad15 | Activation | 0.724 | 2.359 | 0.345 | (31) |

101.	Pan Q, Yang Xh Fau - Cheng YX, Cheng YX. Angiotensin II stimulates MCP-1 production in rat glomerular endothelial cells via NAD(P)H oxidase-dependent nuclear factor-kappa B signaling. (1414-431X (Electronic)).
